# Concurrent Multimodal Data Acquisition During Brain Scanning is within Reach

**DOI:** 10.1101/2021.09.07.459353

**Authors:** Rosa Sola Molina, Gemma Lamp, Laila Hugrass, Russell Beaton, Marten de Man, Lisa Wise, David Crewther, Melvyn Goodale, Sheila Crewther

## Abstract

**Background:** Previous brain-scanning research exploring the neural mechanisms underpinning visuomotor planning and control has mostly been done without simultaneous motion-tracking and eye-tracking. Employing concurrent methodologies would enhance understanding of the brain mechanisms underlying visuomotor integration of cognitive, visual, ocular, and motor aspects of reaching and grasping behaviours. Therefore, this work presents the methods and validation for a high-speed, multimodal and synchronized system to holistically examine neural processes that are involved in visually-guided movement.

**Methods:** The multimodal methods included high speed 3D motion tracking (Qualisys), 2D eye-tracking (SR Research), and magnetoencephalography (MEG; Elekta) that were synchronized to millisecond precision. Previous MRIs were taken to provide improved spatial localization. The methods section describes the system layout and acquisition parameters to achieve multimodal synchronization. Pilot results presented here are preliminary data from a larger study including 29 participants. Using a pincer grip, five people (3 male, 2 female, ages 30-32) reached for and grasped a translucent dowel 50 times, after it was pseudorandomly illuminated. The object illumination was the Go cue. Seven discrete time points (events) throughout the task were chosen for investigation of simultaneous brain, hand and eye activity associated with specific visual (Go cue), oculomotor (1^st^ saccade after Go), motor (Reaction Time; RT, Maximum Velocity: MV, Maximum Grip Width; MGW) or cognitive (Ready, End) mechanisms. Time-frequency analyses were performed on the MEG data sourced from the left precentral gyrus to explore task-related changes time-locked to these chosen events.

**Pilot results:** Basic kinematic parameters including RT, MV, MGW, Movement Time, and Total Time were similar to previous, seminal research by Castiello, Paulignan and Jeannerod, (1991), using a similar task. Although no gaze instructions were given, eye-tracking results indicated volunteers mostly gazed at or near the target object when Ready (72%), and then hardly looked away throughout the rest of the task at the important events sampled here (92% - 98%). At the End event, when lifting the dowel, on average, participants gazed at or near the target object 100% of the time. Although saccades > 100 ms after Go, but prior to RT were made on average in about one fourth (*M* = 13, *SD* = 6) of trials, a mixed model (REML) indicated their latency in timing after the Go was significantly (*F* = 13.376, *p* = .001) associated with RT scores on those trials (*AIC* = 724, *R* _*m*_^2^ = 0.407, *R*_*c*_^2^= 0.420). Neural activity relative to baseline in the beta band was desynchronized for the visually guided reach periods, beginning prior to Go, and remaining sustained until beyond End, after the grasp and lift were executed.

**Conclusion:** This study presents the layout, acquisition parameters and validation for a multimodal, synchronized system designed to record data from the hand, eye and brain simultaneously, with millisecond precision during an ecologically-valid prehension task with physical, 3D objects. The pilot results align with previous research made with single or bimodal data recordings. This multimodal method enables full-brain modelling that can holistically map the precise location and timing of neural activity involved in the visual, oculomotor, motor and cognitive aspects of reach-to-grasp planning and control.

## Introduction

Visuomotor control underpins most actions in the everyday functioning of humans. The neural processes underlying visuomotor control have long been examined using reach-to-grasp (prehension) tasks. During prehension, individual characteristics (e.g. hand size, strength, posture, previous experience) and object properties (e.g. location, size, shape, weight) must be transformed into correct motor coordinates for task execution. Research into the visual processes that drive prehension has shown that processing is enhanced in the visual field region surrounding the hand (Janssen and Scherberger (2015), even when the hand itself is occluded from vision (Perry, Amarasooriya, & Fallah, 2016). Likewise, grasping has been shown to influence object size perception (Bosco, Daniele, & Fattori, 2017), suggesting that prehension also drives visual perception. This may function via an effector-based attention system engaging feedback mechanisms associated with the hand’s location, similar to the visual enhancement that is observed at the end location of an eye movement (Perry & Fallah, 2017). In addition, the planning of saccadic eye movements in parieto-frontal brain areas shifts spatial attention, which improves subsequent visual processing in posterior, visual brain areas via feedback mechanisms (Edwards, Vetter, McGruer, Petro, & Muckli, 2017; Gutteling, van Ettinger-Veenstra, Kenemans, & Neggers, 2010). This highlights that prehension activates neural mechanisms associated with visual attention, oculomotor and manual motor control and visuomotor planning. Thus, hand and eyes are not only driven by these forms of neural activity, but each type of engagement (e.g.: particular movement sequence) also influences neural processing associated with subsequent motor control in unique ways. Consequently, recording eye and hand movements (kinematics) alongside full brain activity is necessary to explore the timing and interactions of all networks involved in visuomotor integration occurring prior to, and throughout, prehensile movements.

To date, a large portion of human brain recordings in visuomotor research have been made in isolation, without motion tracking (Cavina-Pratesi, Goodale, & Culham, 2007; Ehrsson et al., 2000; Gallivan, Cavina-Pratesi, & Culham, 2009; Zaepffel & Brochier, 2012) Culham et al., 2003). When hand movements are recorded during brain scanning, eye-tracking is typically absent (Betti, Zani, Guerra, Castiello, & Sartori, 2018; Bradberry, Gentili, & Contreras-Vidal, 2010; De Sanctis, Tarantino, Straulino, Begliomini, & Castiello, 2013; Di Bono et al., 2017; Mateo et al., 2015; Swett, Contreras-Vidal, Birn, & Braun, 2010; Tscherpel et al., 2020) presumably due to the challenges associated with placement of cameras and lighting, as well as the electrical noise produced by cameras during reach-to-grasp in an fMRI or MEG scanner. Hence, eye-tracking during brain scanning is mostly obtained during reach-to-grasp tasks that lack ecological validity – typically using virtual reality without tactile completion (Committeri et al., 2004; Limanowski, Kirilina, & Blankenburg, 2017), viewing and ‘grasping’ images of objects presented on a screen (Park et al., 2014), or else while viewing the target object via a series of mirrors (Frey, Hansen, & Marchal, 2015; W. Frey et al., 2015; Limanowski et al., 2017). These different methods potentially alter visual and brain processing (Freud et al., 2018).

Thus, detailed hand and finger kinematics during the viewing and grasping of a physical object are seldom measured during brain scanning, and even more seldom recorded synchronously with eye position. Moreover, most research in visually guided prehension focuses on specific topics that rely on one or two recording methods, including:

- Motion-tracking: Factors influencing motor output throughout a reach to grasp, for example, separation of reach and grasp components (Hoff & Arbib, 1993; Jeannerod, 1984), differences between child and adult movements (Zoia et al., 2006), or the time decay of object size vs. object position on reach-to-grasp kinematics (Hesse, Miller, & Buckingham, 2016).
- Eye-tracking: Visual factors influencing visuomotor tasks, for example, target object factors influencing visual awareness (Deplancke, Madelain, & Coello, 2016), or visual attention (Ambrosini & Costantini, 2017), and anticipatory eye fixations (Belardinelli, Stepper, & Butz, 2016), or how eye movement location determines endpoint precision (Ma-Wyatt & McKee, 2007) and eye movement accuracy determines manual interception strategies (Fooken, Yeo, Pai, & Spering, 2016).
- Brain-scanning: Neural activity underlying prehension, for example, the cortical time course for reach to grasp (De Sanctis et al., 2013), the influence of handedness in cortical activation (Begliomini, Nelini, Caria, Grodd, & Castiello, 2008), the neural signatures for action vs. semantic properties of objects (Lee, Huang, Federmeier, & Buxbaum, 2018), or the role of dorsolateral prefrontal cortex in supporting the representation of task-relevant information (Jackson, Feredoes, Rich, Lindner, & Woolgar, 2021).

These methodological divisions result in study designs that support models that are biased towards the modality under consideration, i.e. motor, visual, or cognitive variables, which do not encourage effective testing of holistic theoretical models that attempt to provide a more integrated perspective (Caiani & Ferretti, 2017; Neilson & Neilson, 2004; Schenk, 2010). Indeed, the central question “how is sensory information transformed into purposeful acts?” remains largely unanswered (Milner, 2017).

Hence, we aimed to overcome this challenge by designing a method for concurrent recordings of the eye, hand, fingers, as well as the temporal and spatial brain activations during a reach-to-grasp task. The purpose of this pilot study was to test the multimodal methods developed prior to completing the larger study with the complete data set which will explore the timing and interactions of multiple networks involved in visuomotor integration occurring prior to and throughout prehensile movements of an ecologically valid task. This combination provides the means to develop and test models that provide the full range and temporal sequence of neural activity associated with prehension.

## Methods

### Participants

For the purpose of demonstrating the capabilities of our system, we took a subset of data from a larger MEG study, by selecting participants who had complete MEG, kinematic and eye-tracking datasets for the reach-to-grasp task. Participants included five, dominantly right-handed subjects (two female, three male), aged 30-32 years old, with no reported neuropathological condition. Participants were screened for normal visual acuity, colour vision, stereoacuity, and right hand dominance using a free downloadable iPad app called ‘Eye Test Free-Snellen Chart/Ishihara Test’ (Clark, 1924), the Randot Stereotest (Wang et al., 2010), and Edinburgh Inventory (Oldfield, 1971) respectively. Written, informed consent for participation was given by all participants, and was approved by the La Trobe University Human Research Ethics Committee, HEC16-131. Participants were reimbursed for the time with $50 AUD gift vouchers.

### Apparatus

A diagram illustrating the system developed is presented in Figure 1 below. The system integrated motion tracking of the hand and objects (Fig 1d), eye-tracking of the left eye (Fig. 1c), and full brain MEG (Fig. 1e;) together with a custom-made apparatus (Fig. 1a). The apparatus consisted of a perspex desk that fitted into the MEG chair, with grooves to hold a fibreoptic button box (VPixxTechnologies, 2018), three Perspex objects, and fiberoptic cabling that was connected to an acquisition system controlling the object’s light (Fig. 1b). The acquisition system (VPixxTechnologies, 2018)Fig. 1b) sent TTL signals to the MEG and eye tracker at the blue button press (Fig. 4a), Go (light on; Fig. 4c), RT (button release; Fig. 4d), and light off (2000ms after Go) events for each trial. For every camera frame recorded, the motion tracking system sent a TTL pulse to the MEG acquisition system (Fig. 1d). This allowed millisecond precision alignment of the eye tracking, hand motion and MEG signals with each other for offline processing.

**Figure 1.**
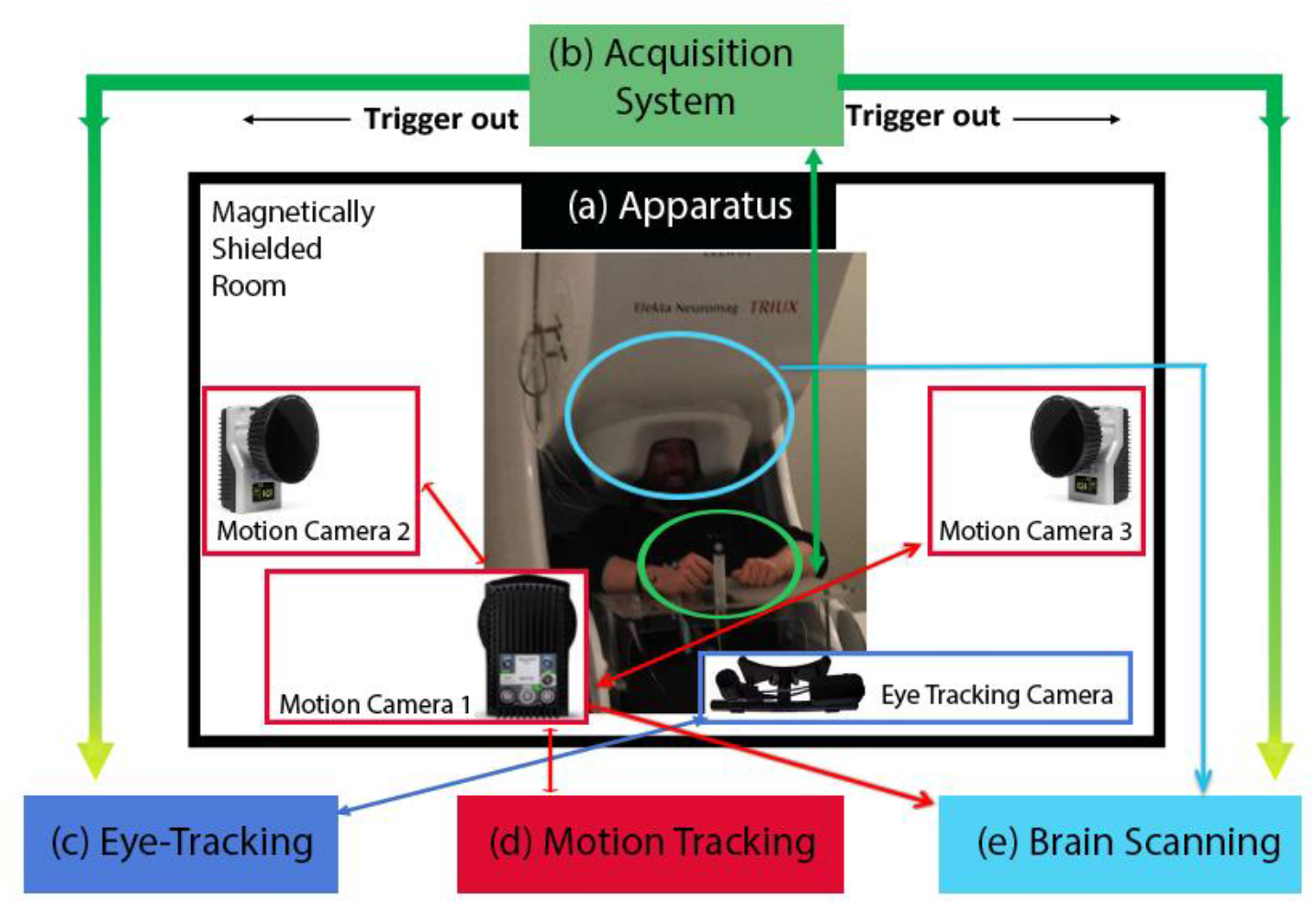
System Designed for Multi-Modal, Full Brain, Synchronized Data Acquisition During an Ecologically Valid Visually Guided Grasping Task Involving Physical Objects. *Note:* Apparatus (a) is controlled by Acquisition System (b) which sends triggers to sync with Eye-Tracking (c) and Brain Scanning (e). Motion Tracking (d) sends triggers to Brain Scanning with every frame (250Hz) recorded to sync with other systems.

### Motion Tracking

To enable optimal recording hand movements in 3D, three infrared motion tracking video cameras (Qualysis, 2018, MRI; sampling rate 250 Hz) were mounted (Qualysis, 2018, Vitek mount) in a magnetically shielded room with two cameras on the walls, and one near the ceiling directly in front of the participant (see Fig. 2a for diagram).

**Figure 2.**
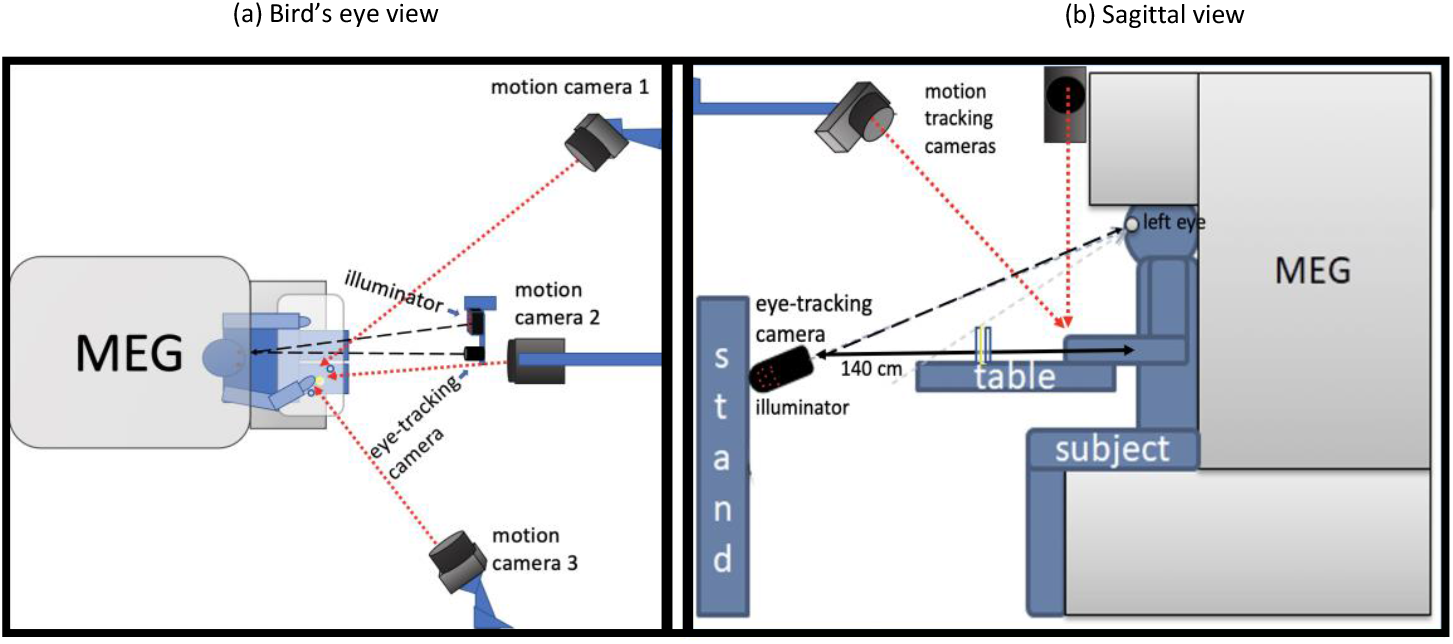
Diagrams for Geometric System Layout in Magnetoencephalography Room. Note: (a) Placement for three motion cameras and eye-tracking camera. (b) Placement of participant’s head and eye-tracking camera height and distance from participant.

The angle of this camera was chosen to provide a direct line of sight to the hand markers throughout the trial. Pilot testing confirmed large camera-induced artefact in MEG recordings was related to receiving event triggers from the acquisition system. Further, connecting the cameras to an analogue acquisition interface placed outside the magnetically shielded room, in order to simultaneously acquire triggers from the acquisition system while video recording produced similarly large artefact in the brain scanning data that could not be filtered. As a result, the motion tracking data could not be synced with the acquisition system whilst online. For this reason, time stamps to mark each video frame were sent directly from the motion tracking computer to the MEG acquisition computer via a BNC cable so that the data could be synced offline. The motion capture system was calibrated to sub mm precision (less than 0.25 mm residuals) by following manufacture recommendations in Qualisys Track Manager software (Version 2018.1, build 4300), using a carbon fibre wand and three spherical markers at constant positions placed on the custom-made desk (Qualisys, 2018).

### Eye-tracking

Monocular gaze position was acquired using a desktop EyeLink 1000 Plus camera (Version 5.15; SR Research Ltd, Ottawa, ON, Canada) with a MEG compatible, long-range mount (SR-Research, 2018). The Eyelink camera processed images via a dedicated laptop, with high-level control through a desktop computer running a custom-made program in Experiment Builder software (version 2.1.140). During recording, the dedicated eye-tracking laptop received TTL signals from the acquisition system (DATAPixx, (VPixxTechnologies, 2018) at four events for each trial (blue button press, Fig. 4a; Go, light on, Fig. 4c; RT, button release, Fig. 4d, and light off, 2000ms after Go) via a modified BCN cable connection with an ExpressCard IEEE 1284 Parallel Adapter card (StarTech.com; SPP/EPP/ECP). Temporal alignment of samples between the two systems was excellent, with root mean square error at RT = 3 ms.

Due to the geometry of the task, standard eye tracking procedures were adapted to ensure the eye was within the trackable range of the camera while not being occluded by the target objects or by the participant’s arm during its reach trajectory as illustrated in Figure 2 and Figure 3b. As the eyes fixated downwards to look at the illuminated dowel, the camera was set lower to the ground compared to screen-based eye-tracking, and the angle was adjusted to capture the greatest surface of the eyes. This camera positioning caused the dowels to partially occlude the right eye from the camera, and made binocular eye tracking impractical, hence we performed monocular gaze tracking of the left eye. The camera and illuminator were mounted on a custom-made wooden stand and placed 140 cm in front of the participant’s midline with the illuminator to the left of the camera. The infrared illuminator was set to the maximum brightness level and its angular position was adjusted until all 20 emitters were directly facing the participant’s left eye. Eye position was calibrated to five points that were marked on the target objects (Fig. 3b). Given that the target objects were towards the edge of the trackable gaze range, calibrations that did not pass the stringent validation requirements of the system were manually accepted. Across participants, the mean calibration errors ranged between 1.61 to 3.46 degrees of visual angle. As discussed below, this limited the degree of spatial precision in our analyses.

**Figure 3.**
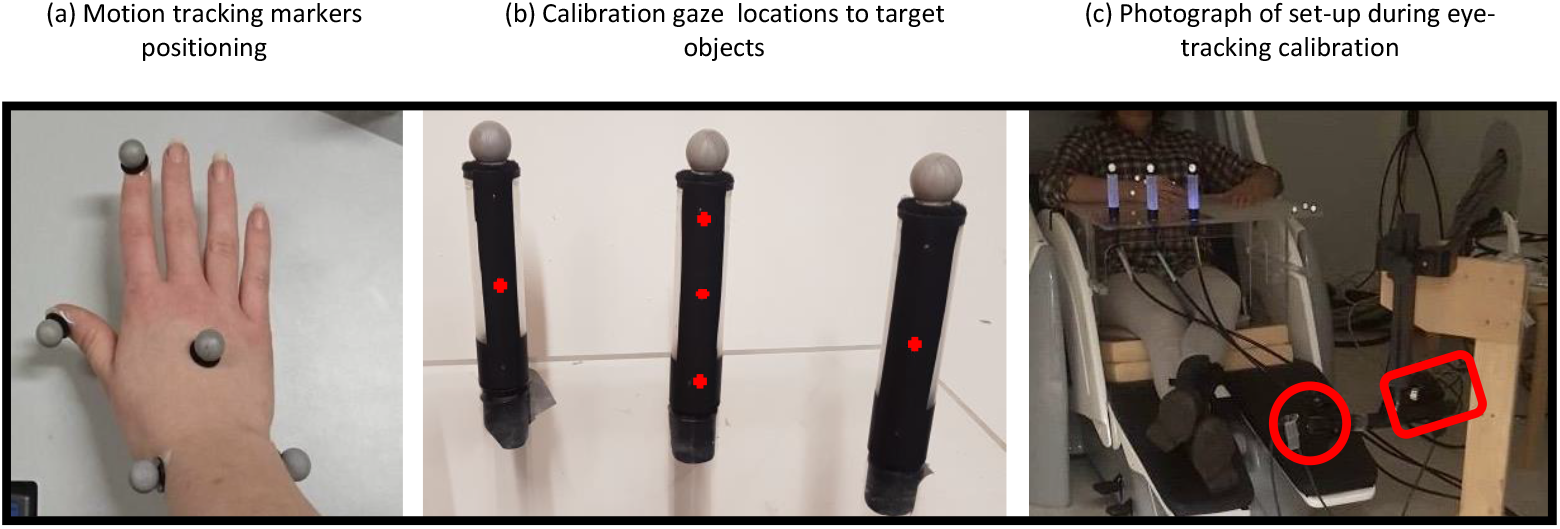
Positioning of Motion Tracking Markers, Eye Tracking Gaze Locations for Calibration and Photograph of Eye Tracking Camera and Illuminated Objects During Eye-Tracking Calibration. Note: (a) Locations named thumb, index, back of hand, medial wrist and lateral wrist; (b) Subject’s view of objects and fixation points (red crosses) during eye-tracking calibration to objects; afterwards, participants rotated objects to view the illuminated side of objects that are visible in panel c. (c) Positioning of objects and eye-tracking camera (red circle) in front and below participant with illuminator (red rectangle) to the left side.

### Magnetoencephalography (MEG) and Magnetic Resonance Imaging (MRI)

MEG data were acquired at the Swinburne Neuroimaging Facility node of the National Imaging Facility, using a 306-channel Elekta Neuromag TRIUX system (ElektaOy, n.d.), with 102 magnetometers and 204 planar gradiometers sampling at 1000 Hz. The MEG was installed in a magnetically shielded room with active flux compensation (MaxShield technology). Temporal alignment of samples between the two systems was excellent (ie., the root mean square error of all samples at RT was = 0.002 s).

Head position was measured using five head position indicator (HPI) coils that were affixed to the participant’s head (on the left and right mastoids and three on the forehead). The positions of the HPI coils, anatomical landmarks (left pre-auricular point, right pre-auricular point, and nasion), and the shape of the participant’s head, were digitized using a 3D Polhemus Fastrack pen. In order to detect blink and cardiac artefact, simultaneous EOG and ECG were recorded with respect to a ground electrode on the left elbow. Vertical EOG was recorded from electrodes that were placed above and below the right eye. ECG electrodes were placed on the left wrist and pectoral muscle. To minimise MEG artefacts from head and shoulder movements throughout the reach-to-grasp task, hardboard foam and cushions were used to adjust the height of the seat so the participant’s head was in contact with the top of the sensor helmet, while their right arm was resting at a 90° angle on the desk. Structural MRI images were obtained using a Siemens 3 Tesla TIM Trio MRI scanner (SiemensMedicalSolutions, n.d.), and used for co-registration of participants’ MEG data with their anatomical data.

### Task & Procedure

The task was based on Castiello, Paulignan, & Jeannerod’s (Castiello, Paulignan, & Jeannerod, 1991) seminal study. Three translucent dowels (2.5 cm diameter and 10 cm height) illuminated by fibre-optic cabling, were placed 35 cm in front of the participant’s hand’s starting point, in front and at 15°, 30°, and 45° from the midline (Fig. 4). A three mm deep cut-out in the table held each of the objects in place. A reflective motion tracking marker (Qualisys, 2018, super-spherical, 19 mm) was affixed to the top of each dowel. Five reflective motion tracking markers (Qualisys, 2018, super-spherical, 9.5 mm) were affixed to subjects’ right thumb, index, back of hand, medial wrist and lateral wrist using double-sided wig tape (Fig. 3a). Once comfortably seated, and after participants’ vision adjusted to the ambient lack of illumination, eye-tracking calibration to pin-sized locations on target objects was performed (Fig. 3b). Afterwards, participants were asked to turn the objects around so that the uncovered side showing complete illumination was visible (Fig. 3c). The grasping task reported here took about five minutes to complete and was the second of seven visuomotor tasks performed in a darkened room.

**Figure 4.**
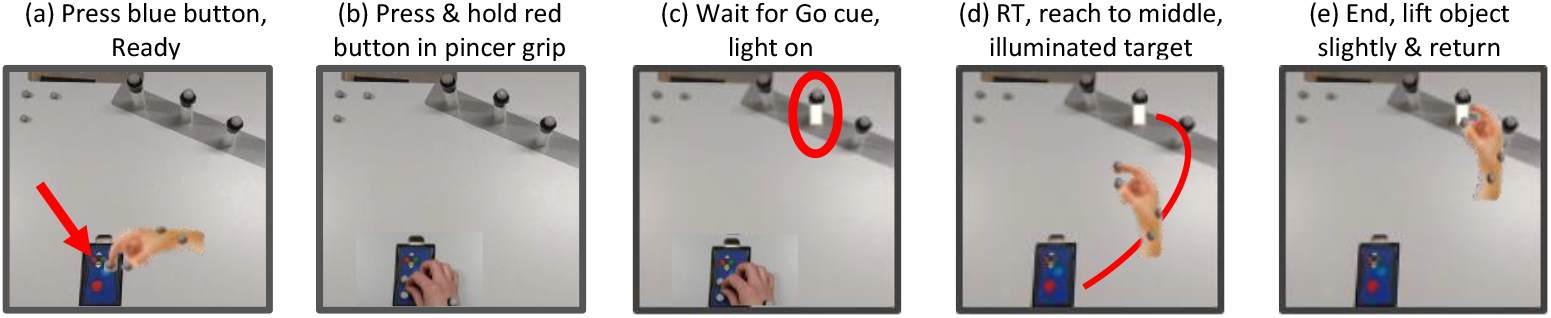
Grasping Apparatus Set-up and Reach-to-grasp Task: Bird’s Eye View of the Three Target Objects with Reflective Markers, and Button Box. *Note:* Task steps: 1) Subjects pressed the blue button on a button box (red arrow), indicating they were ready to begin 2) Subsequently, they placed their index finger and thumb in a pincer grip position while holding down the reverse-coded red button to record release, anticipating the cue. 3) The middle dowel was illuminated pseudorandomly, 2000-3000 ms after the blue button was pressed. 4) A pincer grip was used to reach and lift the dowel as quickly and accurately as possible. 5) Subjects returned the object to its place.

The task included five steps as illustrated in Figure 4. Subjects practised pantomiming the task outside the MEG to confirm they had learnt the sequence, and they were notified that all reaches would be to the center target. Instructions were to grasp as “quickly and accurately as you can”. No instructions were given regarding gaze location. Subjects completed 50, self-paced trials where they reached for and grasped the middle, translucent dowel in a darkened room.

## Data Analysis

To investigate our multimodal system’s capabilities, and demonstrate integration of brain, eye and hand recordings, seven timepoints (events) in reach-to-grasp were chosen that are important in understanding neural processes underpinning cognitive, visual, oculomotor and motor variables in planning and controlling prehension. These seven events were calculated for each trial (see kinematic and eye-tracking data processing).

i. Ready, a time point (one second after the blue button was pressed) was used to illustrate baseline neural activity compared to the other timepoints;
ii. Go, when the light turned on cueing participants to start (i.e. visually evoked neural responses to the object light);
iii. Oculomotor Reaction Time (oRT), the first saccade after Go, but before RT;
iv. Reaction Time (RT), the button release;
v. Maximum Velocity (MV) of medial wrist reflective marker;
vi. Maximum Grip Width (MGW) between index and thumb reflective markers;
vii. End time based on when the object crossed a 5mm. upward threshold

Separate time-frequency analyses were conducted on source-space MEG signals for each timepoint of interest. In addition, the location of eye fixation at each timepoint was investigated. All statistical analyses were performed using IBM SPPS Statistics for Macintosh, IBMCorp (Released 2019).

### Kinematics

Qualisys Track Manager (QTM, Version 2018.1, build 4300, Qualisys) software was used to create a hand model to automatically label the five hand markers according to their anatomical location (Suppl. Fig.1A). Every file was manually examined to ensure the labelling was correct. A custom National Instruments (2018) LabVIEW (Bitter, Mohiuddin, & Nawrocki, 2006) pipeline was written to filter the data (2nd order, low pass Butterworth filter with a cut-off of 125 Hz) and to extract the hand kinematic events: RT, MGW, and End. DATAPixx (VPixxTechnologies, 2018) time stamps were used to identify the Go and RT camera frames. The End of each reach was identified as the camera frame when the z-axis (vertical) position of the spherical marker at the top of the target object exceeded a 5 mm lift threshold. Movement Time (MT) was defined as the latency between End and RT events, and the Total Time (TT) was defined as the latency between the End and Go events. MV was identified based on the medial wrist marker and MGW was identified based on the 3D coordinates of the thumb and index finger. On average, 94% (*SD* = 4%) of the data at each kinematic event was analysed after deletion of data due to participant errors or motion tracking technical errors (Table 2).

### Eye-tracking

DATAPixx timestamps were used to align the Eyelink and MEG data files. Data Viewer (Version 3.2.48, SR Research, 2018) software was used to process eye-tracking data. All data were overlaid onto a 2D photograph of the objects (see Suppl. Figure 3). The objectives of the eye-tracking analyses were to: (i) calculate oRT to the Go cue, and (ii) to determine the gaze position relative to the target object throughout the task, and at precise time points indicating significant kinematic moments. However, it was not practical to determine precise x-y gaze coordinates. Rather, our analyses classified whether fixations and saccades fell within a broad rectangular region of interest surrounding the target object (roughly 25 mm to the left and right of the object and 20mm above and below the object, see Figure 5a). This took account of the calibration errors due to the wide angle recorded previously discussed, and defined whether participants were looking at or near the target object.

**Figure 5.**
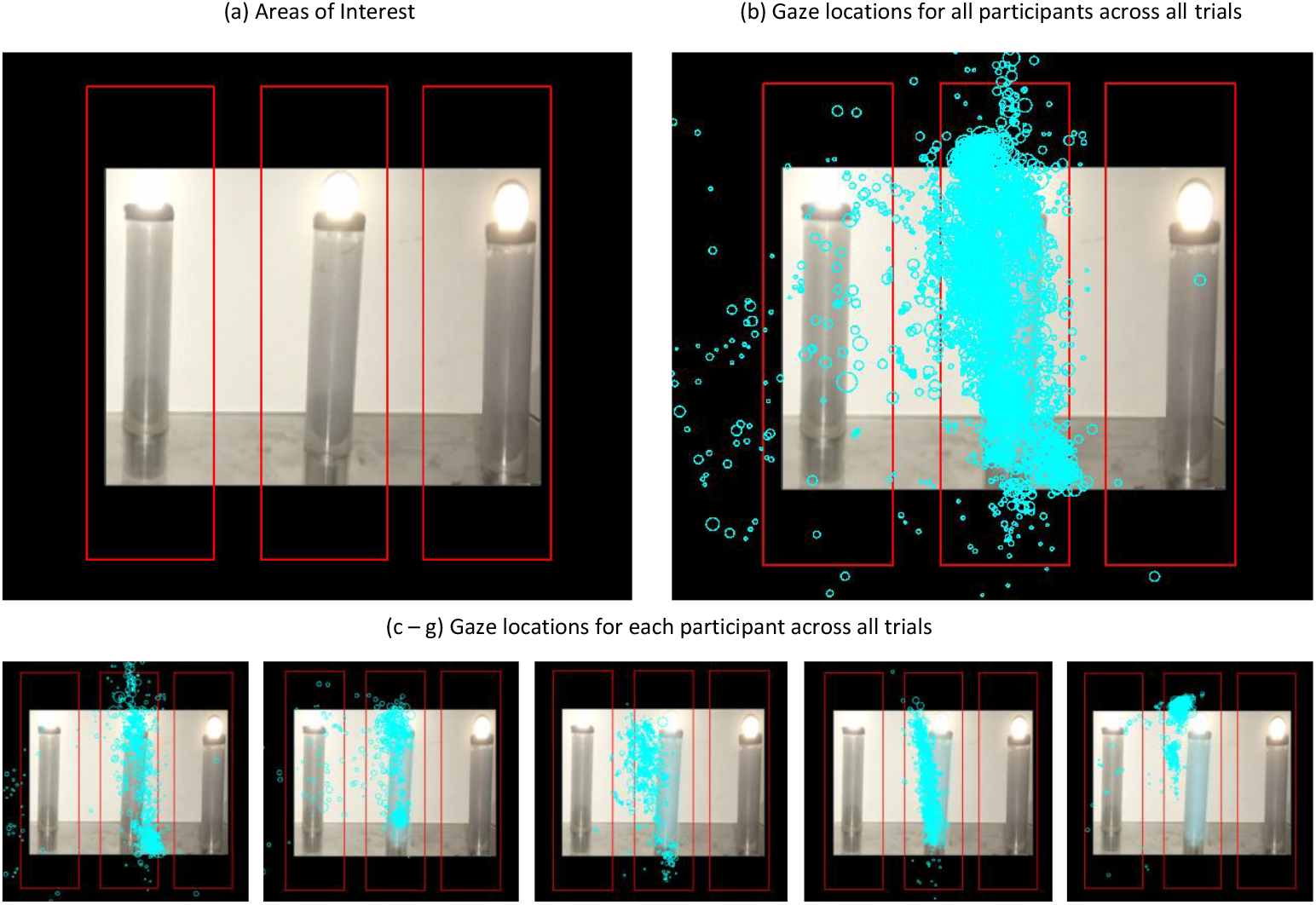
Interest Areas Drawn Around 2D Image of Objects to Define Gaze that is On or Near Target, and All Gaze Locations Recorded During 50 Reach-to-Grasp Trials Together and for Each Participant (n = 5) *Legend*: Blue circles = fixations (gaze location); Larger circles = longer gaze duration.

A Matlab script was developed to extract the relevant eye tracking variables at the time points provided by DATAPixx (Ready, Go, RT) and hand kinematic results (MV, MGW, End) for each trial. Saccades were defined as eye movement speeds that exceeded 22 °/s and resulted in a gaze shift of more than 0.3°. For the purpose of our analyses, we were primarily interested in saccades that occurred during the reach-to-grasp planning period, i.e., the time between Go and RT. The oRT was defined as the onset latency of the first saccade after Go. Express saccades (i.e. those with latencies less than 100 ms) were excluded from the analyses as they are planned using different neural mechanisms, and do not typically correlate with RTs (Fischer, 1986). The number of trials in which a saccade was made during each event was averaged across participants. The location of fixations and saccades at each trial event was calculated as the average number of trials in which fixations and saccades fell within the target object’s area of interest. Oculomotor reaction time occurred in 26% of the trials (first saccade, *SD* = 12%). On average 93% (*SD* = 6%) of the data at each event was analysed after deletion due to eye-tracking or motion tracking technical errors, loss of corneal reflection due to blinks or looking away from the area being tracked (Table 2).

### Magnetic resonance imaging

Cortical reconstruction and volumetric segmentation for the T1-weighed images was performed with the Freesurfer image analysis suite, which is documented and freely available for download (Fischl, 2012). Freesurfer morphometric procedures have been demonstrated to show good test-retest reliability across scanner manufacturers and across field strengths (Fischl, 2012).

### Magnetoencephalography (MEG)

An empty room recording was collected on the day of each recording to capture the noise conditions (with the Qualysis cameras switched on) for subsequent source-modelling. The empty room and task data were pre-processed in a similar way. Temporal Signal Space Separation filtering was applied to each raw recording using MaxFilter software with default settings (Version 2.1, Helsinki, Finland Elekta, 2016). Subsequent analyses were performed using the GUI-based MatLab script Brainstorm (Tadel, Baillet, Mosher, Pantazis, & Leahy, 2011), which is documented and freely available for download online under the GNU general public license (http://neuroimage.usc.edu/brainstorm). MEG data were low-pass filtered (40Hz), down-sampled to 250 Hz, and signal space projection was applied to remove ECG and eye-blink artefact from the task recordings.

Separate event related analyses were conducted for the six chosen timepoints (Go, oRT, RT, MV, MGW, End). Epochs were imported from -3000 to +1000 ms around each event, with additional 1000ms fringes to account for edge-effects. Epochs with unusual activity (e.g.; high amplitude spikes, sensor jumps, unusually high levels of low or high frequency noise) were excluded from subsequent analyses. In addition, trials were excluded if there was a significant Qualisys camera drop out, or if button-release RT were > 500 ms, indicating the participant was distracted. On average, 88% (*SD =* 12%) of the data was included in the MEG analyses for each key task and hand kinematic event. For many trials, there were no saccades detected during the period between the Go cue and RT, hence MEG analyses for this condition included relatively few epochs (*M* = 13, *SD* = 6).

Anatomical scans were aligned with the MEG sensors using six fiducial points (nasion, left and right preauricular points, anterior and posterior commissure and interhemispheric points). An iterative algorithm was used to refine the co-registration based on the additional digitised head points. Source estimation was completed using the default Brainstorm parameters for head modelling with the overlapping spheres technique, and for weighted minimum norm estimate source imaging, with source orientations constrained to the cortical surface. Hilbert transformations were applied to decompose the full cortical source maps for each trial into beta (15-29 Hz) activity and Morlet transformations were applied to create time-frequency maps (3 – 40 Hz) for a region of interest (i.e., scout) in the left precentral gyrus. We used the event-related synchronization/desynchronization method to normalise the time-frequency data relative to baseline. The analyses for the Ready event (i.e. task step 1, blue button press prior to Go) were normalised relative to a 1000 to -4 ms baseline. The Ready event initiated a 2000 – 3000 ms waiting period, during which the participant held down the red button until the light cued them to perform a reach. As illustrated in Figure 6, pre-motor beta was elevated during the waiting period across participants. Hence, this provided a stable baseline for analyses of the subsequent task events. The Go and oRT analyses were normalised with respect to a -1500 to -1000 ms baseline. The RT, MV, MGW and End analyses were normalised relative to a -2000 to -1500 ms baseline.

**Figure 6.**
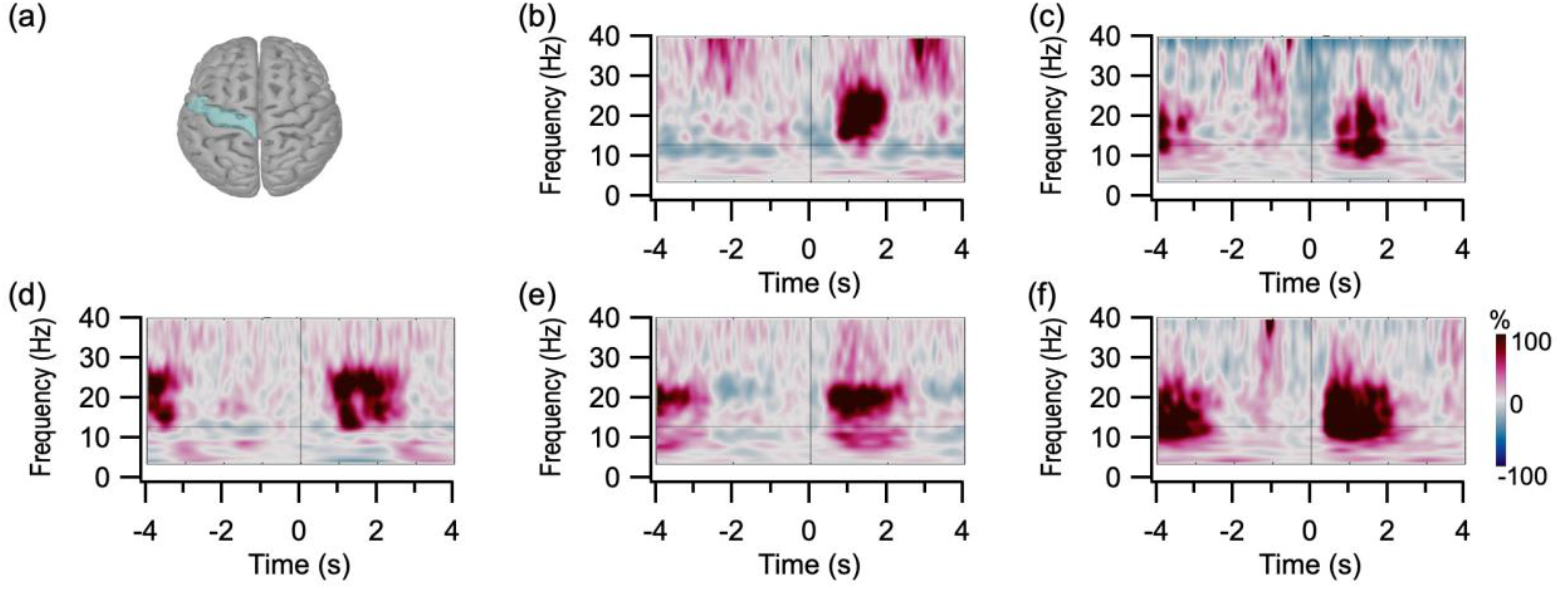
Changes in Beta Synchrony After the Ready Event are Illustrated for a Region of Interest in the Left Precentral Gyrus (a) for Each Participant (b-f) (n = 5). *Note:* Morlet time-frequency plots (3 – 40 Hz) for the x-axes are centered on the moment in time when the participant pressed the button blue button to indicate they were ready start the trial. The time frequency plots were normalized relative to a -1000 to -4ms baseline, with the magenta and blue colors representing relative increases and decreases in synchronous activity respectively.

## Results

### Hand, Eye & Brain

The results presented in Figure 7 illustrate the estimated hand positions associated with group means for kinematic events (Fig. 7a) and location of eye gaze (Fig. 7b) alongside the group average cortical maps (Fig. 7c) of beta band synchrony (15-30 Hz) for the reach-to-grasp time points of interest. The activity across the left hemisphere motor and premotor cortices indicates strong motor synchronization prior to the Go cue during the waiting period (red in Figure 7c). Further, in comparison to baseline, the extended but weaker beta desynchronization can be observed throughout the entire reach-to-grasp trial, at Go, oRT, RT, MV, MGW and End (blue in Figure 7c).

**Figure 7.**
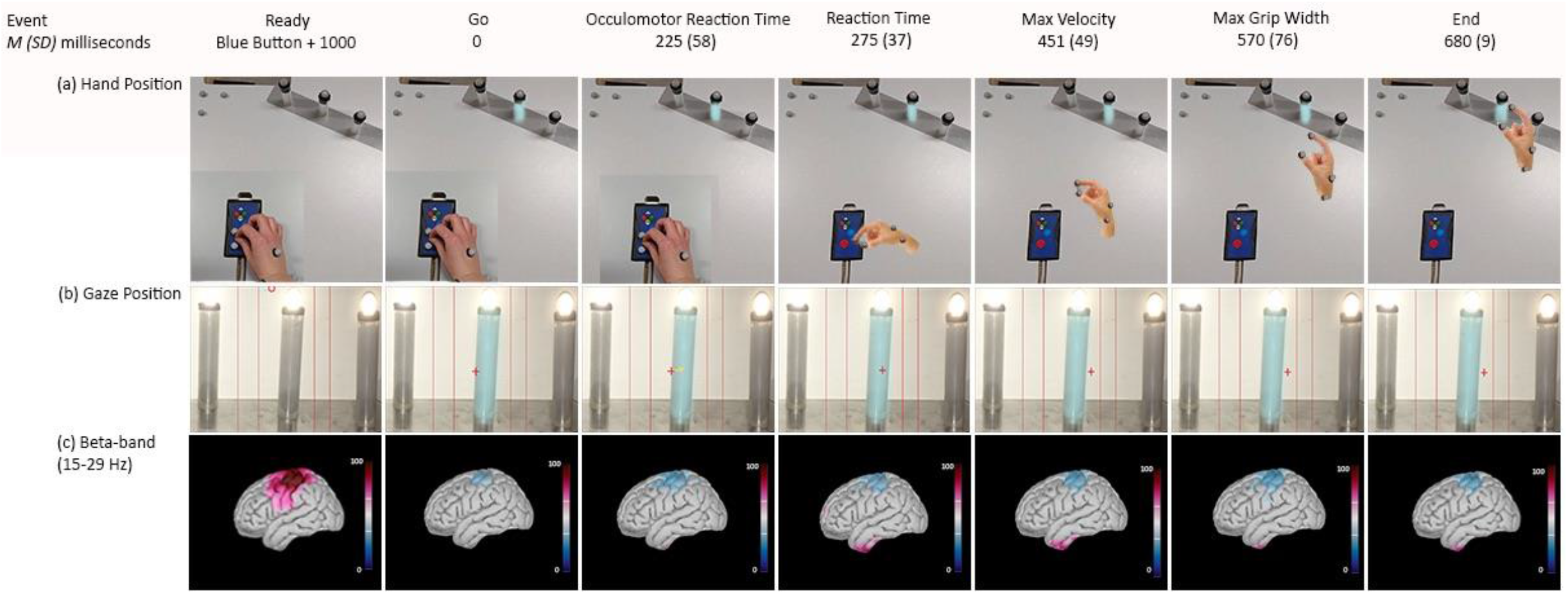
Multimodal Illustration of Hand, Eye and Brain Activity During the Reach-to-Grasp Task for Events of Interest (n = 5): Ready, Go, oRT, RT, MV, MGW, and End. *Note:* Event statistics represent group averaged kinematic and oculomotor results. (a) Estimated hand position at each event of interest for one participant across one trial. (b) Gaze location (red cross) at each event overlayed on an image of target object for one participant (no. 077) across one trial (no14). (c) Group averaged cortical maps of event-related changes in beta power (15-30 Hz Hilbert transform), with magenta indicating synchrony and blue indicating desynchrony, relative to the baseline periods for each event (see Figure 6).

### Kinematics

Descriptive statistics for the four kinematic time points of interest are displayed in Table 1. Across participants, the average TT (from Go to End) was 691 ms (*SD =* 123 ms) and the average MV was 829 mm/s (*SD =* 167 mm/s).

**Table 1.** Descriptive Statistics and Percentage of Movement Time for Reaction Time, Max Velocity, Max Grip Width, and End for 50 Reach-to-Grasp Trials and n = 5

| Variable name | $M$ ($SD$) trials included<br>per subject | $M$ ($SD$) time after Go<br>in milliseconds | Percentage of<br>Movement time |
| --- | --- | --- | --- |
| Reaction Time (Button release) | 47 (2) | 275 (37) | 0% |
| Max Velocity | 46 (4) | 176 (49) | 43% |
| Max Grip Width | 47 (1) | 297 (76) | 73% |
| End (Object Lift) | 48 (1) | 405 (9) | 100% |

*Note:* Movement Time = RT to End.

### Eye-tracking

Descriptive statistics for the seven, eye-tracking time points of interest are displayed in Table 2 below. Overall visualization of the data across participants showed the large majority of fixations throughout the block of trials were on or near the target (See Figure 5). Several saccades made to the left target and beyond occurred mostly between trials, and only one saccade was made to the right target interest area. Persistent fixation on target was likely due to participants’ prior knowledge of the task being limited to the center target. Additionally, participants heads were immobilized (see Methods) and aligned to their midline, while the objects were placed to the right, at an angle. This made saccades to the object farthest on the right a challenging stretch for the eye muscles.

**Table 2.** Eye-tracking Descriptive Statistics for timepoints Ready, Go, Oculomotor Reaction Time, Reaction Time, Max Velocity Time, Max Grip Width Time, and End: Means and Standard Deviations and Percentage of Trials with a Saccade at Timepoint, the average percentage of trials with fixations on target, average percentage of trials with saccades on target, and average for n = 5.

| Timepoint | *M (SD)* trials included per subject | *M (SD)* % trials with saccade at timepoint | *M (SD)* % fixation on target | *M (SD)* % saccade on target |
| --- | --- | --- | --- | --- |
| Ready | 44 (7) | 20% (18%) | 72% (22%) | 70% (19%) |
| Go (Light On) | 50 (0) | 14% (18%) | 95% (9%) | 83% (20%) |
| Oculomotor Reaction Time | 13 (6) | 26% (12%) | n/a | 100% |
| Reaction Time (Button Release) | 47 (2) | 23% (26%) | 95% (12) | 93% (16%) |
| Max Velocity | 45 (4) | 18% (24%) | 92% (18%) | 94% (13%) |
| Max Grip Width | 45 (4) | 22% (27%) | 98% (5%) | 94% (14%) |
| End (Object Lift) | 48 (1) | 27% (23%) | 100% (1%) | 91% (16%) |

Note: Ready = timepoint at blue button press +1000 ms. Trials and percentages are rounded to the nearest whole number. Large *SD*'s reported at fixations/saccades on target during Reaction Time, Max Velocity, Max Grip Width, and End time are due to four out of five subjects fixating about 100% on target.

Throughout the reach-to-grasp, beginning at Go, participants mostly gazed on the target object, eliminating the need to make saccades towards the target after the Go cue (Table 2). Indeed, few trials included a valid saccade beginning >100 ms after Go, but prior to button release. This pattern in our data is likely associated with participants’ training to establish prior knowledge of the task and location of the target object (Adam, Buetti, & Kerzel, 2012). Nevertheless, as the first saccade after a visual cue in prehension tasks is known to incorporate motor and attentional neural processes, it was included as a time point of interest for MEG analysis. Mean time to first saccade after Go was 225 ms (*SD* = 58). This is about 50 ms. before average motor RT (*M* =275 ms).

### Hand and Eye Correlations

Throughout the task, oRT was positively correlated with RT as displayed in Figure 8. A mixed model (Thejamoviproject, 2021), including Time (trial number) as a fixed factor, Participant ID as a random factor and oRT as a covariate, demonstrated that oRT significantly contributed (*F* = 13.376, *p =* .001) to the model, explaining 40% (*R*_*m*_^*2*^ = 0.407) of the variance in RT scores associated with Time (*F* = 0.770, *p* = 0.771). Without oRT, fixed task conditions (Time) explained 10% (*R*_*m*_^*2*^ = 0.099, *F* = 0.862, *p* = 0.724) of the variance in RT scores. Furthermore, random individual factors (Participant ID) associated with variance in RT scores across trials dropped from 35% (*R*_*c*_^*2*^= 0.448) to 1% (*R*_*c*_^*2*^= 0.420). This greatly strengthened the model (*AIC* = 724 compared to *AIC* = 2359 without RT). Similar results were found when running an equivalent model with RT as a covariate explaining variance across oRT scores.

**Figure 8.**
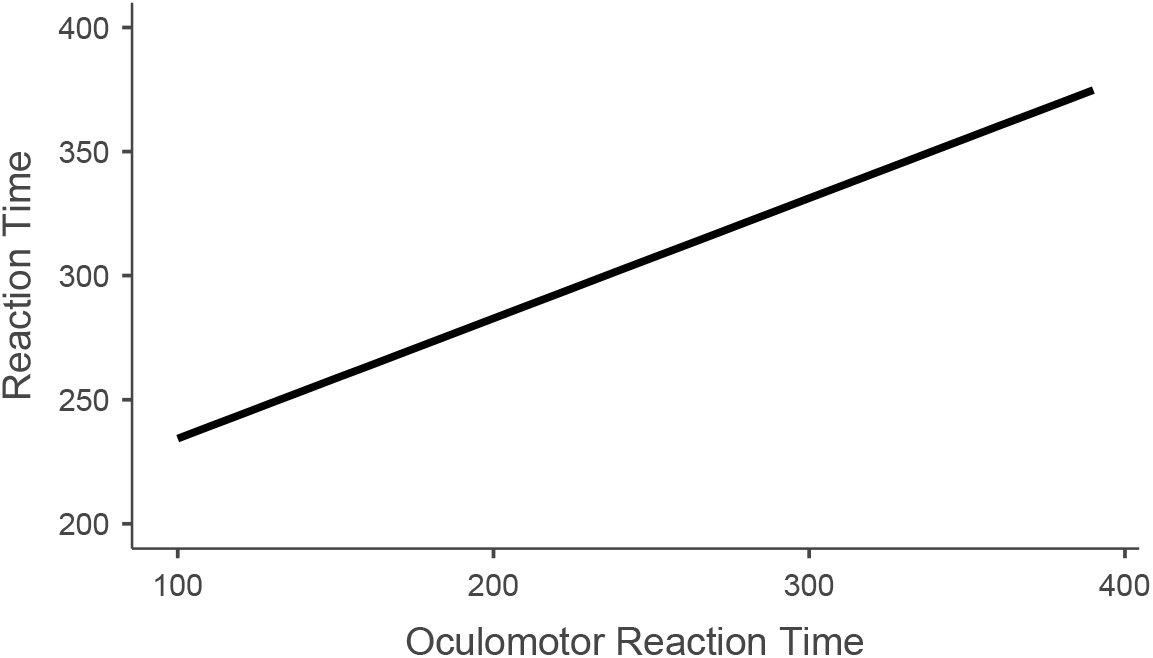
Effects Plot from a Mixed Model (REML) with Time (Trial No) as a Fixed Factor, Participants as a Random Factor and Oculomotor Reaction Time as a Covariate, Illustrating the Relationship Between Oculomotor Reaction Time and the Hand’s Reaction Time Scores Across Time (1-50 Trials) and Participants (n = 5) *Note:* This image includes 72 trials during which participants made a saccade ≥ 100 ms after the Go cue and prior to RT.

### Magnetoencephalography

In order to illustrate the changes in brain activity for different time points during the visually guided reach-to-grasp trials, a set of time-frequency analyses were performed on the MEG data. As illustrated in Figure 7c, there were robust task-related changes in beta synchrony over the motor cortex. The time-frequency maps of the left precentral gyrus (area = 32.84 cm^2^) (Fig. 9, left panels) show that beta band activity was desynchronized for the visually guided reach periods (Fig. 9b–g), from Go to End, relative to the anticipation (Ready) period shown in Figure 9a. Group average plots of the event-related changes in beta synchrony (Fig. 9, 15-30Hz black traces on the right panels), show that desynchronization tended to commence even prior to the Go cue (Fig. 9b), and was sustained beyond when the grasp and lift were executed (Figure 9g). Although there were individual variations in the strengths of these effects, this overall pattern of beta band activity was highly consistent across participants (Fig. 9, grey traces on right panels).

**Figure 9.**
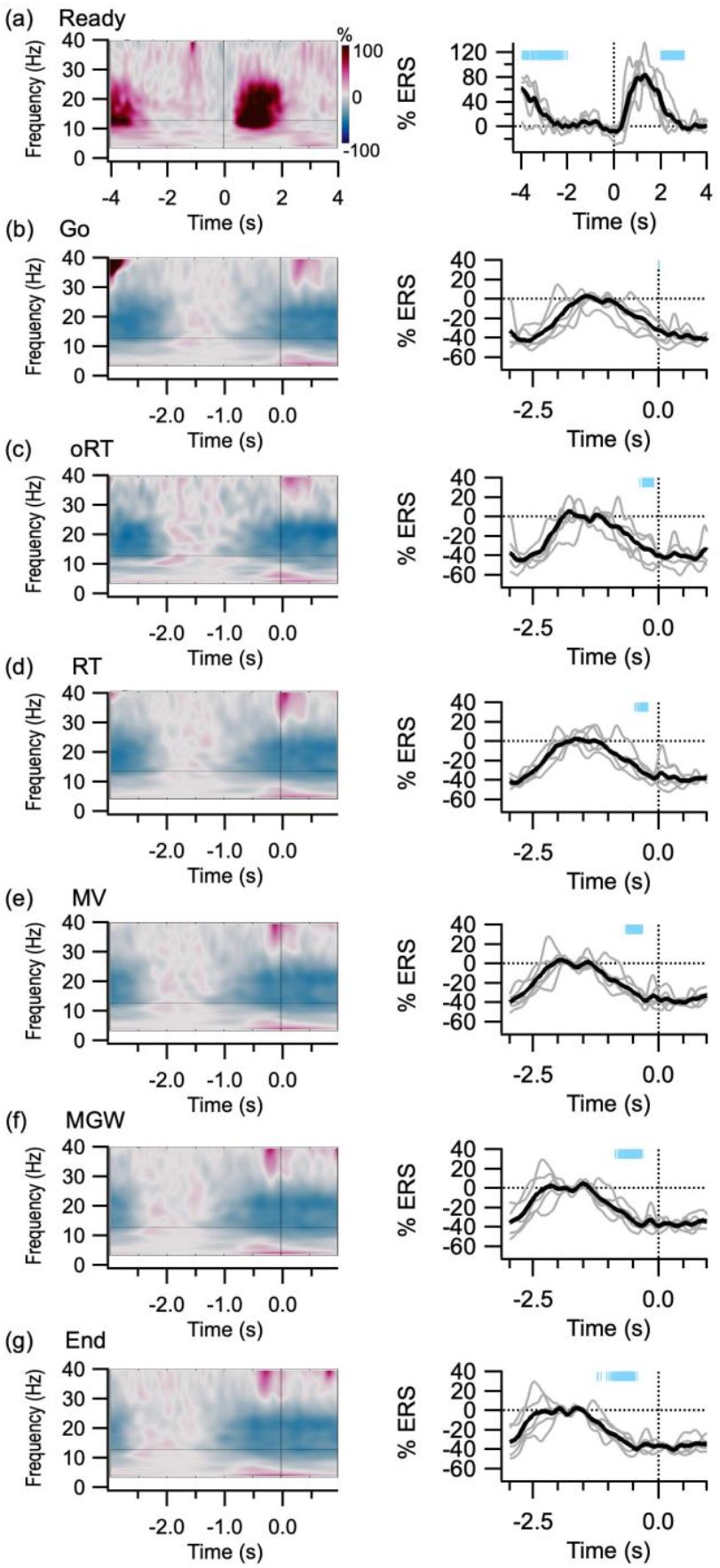
Time Frequency Results (n = 5) for the Left-Precentral Region of Interest at Key Events Throughout the Reach-to-Grasp Task. *Note:* For the pre-trial Ready analyses (a), baseline normalisation was performed relative to the 1s period prior to the blue button press. For other key events, baseline normalisation was performed relative to a period when participants were waiting for the trial to start, i.e.; -1.5 to -1 s prior to the Go cue (b) and oRT (c) events, and -2 to -1.5 seconds prior to the motor RT (d), MV (e), MGW (f) and End (g) events. The time-frequency maps in left panels illustrate sustained changes in beta band activity relative to the baseline periods (Morlet transform, 3 – 40Hz). The magenta and blue colours represent changes in synchronisation and desynchronization respectively. The line plots on the right illustrate changes in beta activity (15-30Hz, Hilbert transform) surrounding each event (t = 0 on the x axes), with the group average in black, and individual traces in grey. The cyan markers at the top of each plot illustrate the distribution of Go event times relative to the other time points.

## Discussion

We have developed and validated a system that records concurrent hand, eye and brain activity with superior spatio-temporal resolution during a reach to grasp movement. Results presented here provide evidence for the system’s robustness and synchrony in sampling. To our knowledge, this is the first time that concurrent eye, hand and full-brain activity have been recorded with high temporal resolution during an ecologically valid task with physical, 3D objects. Thus, comprehensive multimodal models of the brain and body processes involved in prehensile planning and performance can now be tested. Furthermore, the time course of interactions between known visual, visuomotor, motor, and fronto-parietal networks can now be established and extended to include known subcortical and cerebellar structures. These methods offer solutions for current limitations in motor research by eliminating existing biases that occur due to drawing conclusions from only one or two neurophysiological modalities.

Our kinematic results confirm that hands can be successfully tracked in three dimensions from within a magnetically shielded room concurrently with the acquisition of MEG brain signals and eye tracking. Shorter RTs and MTs, as well as an earlier MVs indicate our task was less challenging than the original task replicated here (Castiello et al., 1991). Specifically, the shorter time to MGW that was reached later on in the movement (73% vs. 61%) identifies the grasping component was simpler than in the original task (Paulignan, Jeannerod, MacKenzie, & Marteniuk, 1991b). In contrast, the percentage of MT taken to reach MV, known as the reaching component, was similar to original results (Castiello et al., 1991; Paulignan, MacKenzie, Marteniuk, & Jeannerod, 1991a). This kinematic pattern suggests the reaching component of the movement was similar to original research and was not greatly influenced by the task set-up, but rather the grasping component associated with the object size or positioning. As our task involved a larger target, and a greater distance between the target object and two flanking targets when compared to the original task, easier grasp formation was apparent and likely facilitated faster prehensile planning.

Successful eye-tracking indicated a measure of overt attention and timing of eye-movements during, or leading up to, any point of interest during basic reach-to-grasp tasks in MEG can be recorded with millisecond precision alongside motion tracking. Although participants were not given instructions regarding gaze location in our first series of experiments in order to provide ecological results, overall, they fixated on or near the target prior to and during the reach and grasp. Consistent gazing on target suggests that participants were acquiring visual cues of the object to aid in grip formation prior to and throughout the movement, as well as visual feedback of the approaching fingers to ensure a successful grasp in the later part of the movement (Brouwer, Franz, & Gegenfurtner, 2009). These multimodal methods will permit researchers to test hypotheses regarding the neural mechanisms for increased visual processing close to the hand during prehension (Perry & Fallah, 2017).

Similar to previous research in which the object appeared in the location where participants were already looking (Adam, Buetti, & Kerzel, 2012), few trials contained a valid saccade in response to the Go cue (i.e., light turning on). Nevertheless, when saccades were made, their latency to the Go cue (oRT) was associated with motor response latency of the hands (RT), supporting the notion of tight neural coupling for the initiation of eye and arm movement (Gribble, Everling, Ford, & Mattar, 2002; Land & Tatler, 2009). Additionally, the average 50 ms. oRT preceding motor RT falls within the 40-100 ms. window previously reported in visuomotor tasks (Angel, Alston, & Garland, 1970; Prablanc, Echallier, Komilis, & Jeannerod, 1979; Suzuki, Izawa, Takahashi, & Yamazaki, 2008). These MEG results support previous findings indicating beta band activity over motor and somatosensory cortex and show a clear desynchronization of up to two seconds leading up to and throughout movement (Amengual, Marco-Pallares, Grau, Munte, & Rodriguez-Fornells, 2014; Barratt et al., 2017; Fairhall, Kirk, & Hamm, 2007; Heinrichs-Graham, Arpin, & Wilson, 2016; Jochumsen et al., 2017), and a strong resynchronization beginning around 0.5-1 second after movement is finalised (Ready), when holding a stationary position in expectation of a following motor command or movement (Amengual et al., 2014; Barratt et al., 2017; Fairhall et al., 2007; Gilbertson et al., 2005). Overall, the timing of the largest peak in beta band desynchrony appears to fall somewhere between the initial eye and hand motor response. Individual differences in the timing of the beta desynchrony noted here have been previously reported (Jochumsen et al., 2017) and found to contain a significant genetic factor (Becker et al., 2018).

As the MEG results presented here denote changes in neural activity relative to baseline neural activity, it is important to consider the baseline used, as this can alter interpretation of the findings (Camacho, Quiñones-Camacho, & Perlman, 2020). Our baseline was taken during a time window when participants were using a precision grip to hold down the red button, in anticipation of the light cue. Thus, this task baseline likely reflects motor control and associated visually driven attentive processes. This indicates our results are not reporting a change of neural activity compared to ‘resting cognition’, but rather a change compared to an already dynamic, cognitively driven process, including somatosensory feedback of the button being pressed, and attentiveness related to imminent motor response planning. Therefore, results calculated during specific timepoints of the reach-to-grasp may not represent the full degree of neural change occurring during these stages of the task when compared to a baseline that is completely unrelated to the task. In effect, the results reported here highlight neural activity underlying *visually controlled, online* prehension, minus any ‘offline’ memory-based motor control. Future investigations may include a timed gap between trials during which participants place their hand on the table, to use as a baseline that does not involve a motor component. Ultimately, these MEG results illustrate the robust nature of motor related beta activity over somatosensory and motor cortex, and highlight the need to explore more specific neural signatures associated with diverse kinematics.

The design of these methods improves previous methodologies used in visuomotor brain mapping by enabling the impartial exploration of neural activity associated with visual, motor, visuomotor, or oculomotor processes with the highest spatiotemporal resolution currently available. This will provide a more accurate multimodal view of grasping processes (Betti, Castiello, & Begliomini, 2021). Subsequently, associated brain analysis may classify neural signatures into those more related to visual properties, hand movements, or eye movements, and outline which factors influence their individual contributions and interactions (Guo, Nestor, Nemrodov, Frost, & Niemeier, 2019). Overall, these methods improve previous visuomotor research in three main ways.

Firstly, including motion cameras that have high temporal alignment with brain scanning enables improved disambiguation of the visually-driven, feedforward mechanisms and visually-driven feedbackwards mechanisms occurring throughout prehension when new objects visually appear or are perturbed (Scott, 2016). This is possible due to the increased temporal specificity acquired by time-locking neural analyses to the moment the hand-in-motion begins to deviate in response to an object appearance or change in location, rather than time-locking analysis to the time point the object changes. Additionally, the top-down, feedback effects of oculomotor and motor planning that occur during visually guided grasping movements are known to influence visual processing of location and space differently within the two visual streams (Lehky & Sereno, 2019). Thus, the high temporal resolution provided by concurrent, synchronized motion tracking and eye tracking permits identifying the time period(s) leading up to a change in hand or eye movement that is associated with top-down motor planning in order to temporally locate neural activity associated with feedback effects in visual processing that is associated with subsequent motor patterns. Comparing visual processing that includes motor planning feedback effects in the two visual streams and to visual processing during other visually-guided behavior will permit a complete mapping of visual processing associated with grasping. We envision these multimodal methods will aid in answering longstanding questions regarding which locations across the two visual streams -and at what time(s) relative to the visual onset-, parallel or interactive neural processing occurs during prehension planning and online motor control (Ferretti, 2021; Milner, 2017; van Polanen & Davare, 2015).

Secondly, including eye-tracking alongside motion cameras during brain scanning enables disambiguating the motor planning processes of the hand from those of the eyes. This is possible by time-locking neural analyses to the moment both hand and eye movements begin rather than ignoring or interpreting concurrently increased neural activity leading up to saccades in networks known to plan eye movements as part of the motor planning underlying hand movements. Further, this permits exploration of the feedback provided by oculomotor attention mechanisms that enhance visual processing at the end of an eye movement in addition to potential visuomotor attention mechanisms enhancing visual processing around the hand (Perry et al., 2016; Perry & Fallah, 2017). Additionally, synchronized multimodal recording permits exploration of the extent to which extrinsic (e.g.: object shape, location or crowding), intrinsic (e.g.: purpose and prior knowledge of object) and cognitive factors (e.g.: goals, task) influence prehensile kinematics (Egmose & Koppe, 2018) and relate to visuomotor (Cavina-Pratesi et al., 2018; Takahashi et al., 2017) and/or oculomotor neural processing under various conditions.

Finally, including synchronised, high frequency sampling and MEG technology permits exploring neural activity associated with various strategies for successful motor control, learning and types of errors known to be associated with individual neural differences (Tomassini et al., 2011). This is possible by running multivariate analysis correlating individual participants’ trial-by-trial fluctuations in neural activity associated with specific kinematics (Gu, Wood, Gribble, & Corneil, 2016) and eye-movements. Although similar explorations have been conducted, the current methods improve spatial resolution previously obtained with combined EEG & motor tracking (Amengual et al., 2014; Guo et al., 2019; Sburlea & Muller-Putz, 2018), and temporal resolution obtained with combined fMRI and motor tracking (Budisavljevic et al., 2017; Di Bono et al., 2017; Filimon, Nelson, Huang, & Sereno, 2009). Furthermore, correlating individual neural signature variations to individual kinematic profiles will provide an intrinsic link between neural population activity and specific aspects of motor control that is currently lacking (Lehky & Sereno, 2019). This could help distinguish individual from collective neural signatures, and may clarify to what extent visual, visuomotor, motor or oculomotor aspects are responsible for the motor deficits apparent in stroke or developmental and neurodegenerative conditions.

Together, these improvements in brain mapping might provide more specific targets for clinical studies developing rehabilitation programs. Indeed, with correct camera positioning, a wide range of actions can now be recorded during brain scanning that might further our understanding of stroke lesions, developmental disorders, motor diseases, and the neural processing associated with motor impairment observed in most neural disorders. This would potentially lead to improved rehabilitation protocols that delineate both visual and motor behaviours associated with promoting healthy visuomotor functioning. Further, these multimodal methods offer a means to further develop technological applications using brain-controlled robotic devices aiming to restore autonomy in individuals with spinal injuries resulting in tetraplegia and paraplegia (Ajiboye et al., 2017; Soekadar et al., 2016).

When setting up the current system, the main difficulty encountered concerned the positioning of the eye tracking camera such that the eye was not occluded by the target object or the hand during the reach-to-grasp task. This resulted in a reduction in the spatial accuracy of eye-tracking signals. Thus, the visual behaviour reported here was limited to a general estimation of when the eyes were focused on or near the target object rather than a precise 3D spatial location. Consequently, we were unable to determine whether fixations prior to grasping focused on where the index finger (first digit to touch) or thumb (associated with subsequent object manipulation) was going to make contact (Belardinelli et al., 2016; Betti et al., 2018; Cavina-Pratesi & Hesse, 2013; Voudouris, Smeets, Fiehler, & Brenner, 2018), or else on the centre of target mass, associated with prior knowledge of the task (Voudouris, Broda, & Fiehler, 2019). Therefore, the eye-tracking camera positioning used in these methods is limited. Future improved investigations might explore using existing eye-tracking camera systems that are linked to a hot mirror for calibration that is placed directly in front of vision, through which participants can naturally view target objects (SR-Research, 2018). This would permit calibration to a wider visual angle, enable binocular eye-tracking, and allow any object to be placed directly in front and on both sides of the participant. This would increase ecological validity, improve spatial accuracy, and provide the 3D gaze position information lacking in this study (Bosco, Breveglieri, Hadjidimitrakis, Galletti, & Fattori, 2016).

Several additions might further improve these methods. The inclusion of fMRI to locate early visual cortical and subcortical regions known to process specific visual stimuli would improve spatiotemporal mapping of the time course of visual processing associated with subsequent visuomotor transformations. In addition, future investigations seeking to explore the involvement of somatosensory suppression associated with goal-directed visually guided movement and grip formation may include pressure sensors on the target objects, in order to increase temporal specificity of the tactile stimulus received, and locate the associated neural response (Voudouris et al., 2019). Moreover, future studies may want to include electromyography alongside the modalities included here, in order to identify a more precise order of the timing of motor signals involved in reach to grasp planning and control (Betti et al., 2018; Gribble et al., 2002; Sburlea & Muller-Putz, 2018).

In conclusion, the methods validated here propose a solution for current methodological limitations that have resulted in a fragmented understanding of the cortical underpinning of reach-to-grasp movements. Previously, this research has discussed limited neural networks, or else specific timepoints or frequencies in limited networks and how they are associated with particular aspects of prehension. In response, the synchronised multimodal system presented here records from the hand, eye and brain. These methods enable the simultaneous and unbiased investigation of attentive, visual, motor, oculomotor and cognitive aspects of reach to grasp movements that have previously been explored in isolation or pairs. Results validating the system replicate previous findings in motor, visual, oculomotor, and neural research, evidencing the system’s robustness and synchrony in sampling. Future investigations using these methods and improved eye-tracking may investigate deep brain structures and cerebellum alongside the cortices, allow simultaneous investigations into specific hand and gaze movement patterns, and outline the complete time course and interaction for known cortical visual networks associated with reach-to-grasp movements. Ultimately, this multimodal method enables full-brain modelling that can holistically map the precise location and timing of all neural activity involved in reach-to-grasp planning and movements.

